# State-aware detection of sensory stimuli in the cortex of the awake mouse

**DOI:** 10.1101/499269

**Authors:** Audrey Sederberg, Aurélie Pala, He Zheng, Biyu J. He, Garrett B Stanley

## Abstract

Cortical responses to sensory inputs vary across repeated presentations of identical stimuli, but how this trial-to-trial variability impacts detection of sensory inputs is not fully understood. Using multi-channel local field potential (LFP) recordings in primary somatosensory cortex (S1) of the awake mouse, we optimized a data-driven cortical state classifier to predict single-trial sensory-evoked responses, based on features of the spontaneous, ongoing LFP recorded across cortical layers. Our findings show that, by utilizing an ongoing prediction of the sensory response generated by this state classifier, an ideal observer improves overall detection accuracy and generates robust detection of sensory inputs across various states of ongoing cortical activity in the awake brain, which could have implications for variability in the performance of detection tasks across brain states.

**Author summary:** Establishing the link between neural activity and behavior is a central goal of neuroscience. One context in which to examine this link is in a sensory detection task, in which an animal is trained to report the presence of a barely perceptible sensory stimulus. In such tasks, both sensory responses in the brain and behavioral responses are highly variable. A simple hypothesis, originating in signal detection theory, is that perceived inputs generate neural activity that cross some threshold for detection. According to this hypothesis, sensory response variability would predict behavioral variability, but previous studies have not born out this prediction. Further complicating the picture, sensory response variability is partially dependent on the ongoing state of cortical activity, and we wondered whether this could resolve the mismatch between response variability and behavioral variability. Here, we use a computational approach to study an adaptive observer that utilizes an ongoing prediction of sensory responsiveness to detect sensory inputs. This observer has higher overall accuracy than the standard ideal observer. Moreover, because of the adaptation, the observer breaks the direct link between neural and behavioral variability, which could resolve discrepancies arising in past studies. We suggest new experiments to test our theory.

## Introduction

The large majority of what we know about sensory cortex has been learned by averaging the response of individual neurons or groups of neurons across repeated presentations of sensory stimuli. However, multiple studies in the last three decades have clearly demonstrated that sensory-evoked activity in primary cortical areas varies across repeated presentations of a stimulus, particularly when the sensory stimulus is weak or near the threshold for sensory perception (1-3), and have suggested that this is an equally important aspect of sensory coding as the average response (4-6). Variability is thought to arise from a complex network-level interaction between sensory-driven synaptic inputs and ongoing cortical activity, and single-trial response variability is partially predictable from the ongoing activity at the time of stimulation. A large body of work has focused on characterizing this relationship between notions of cortical “state” and sensory-evoked responses (7-13), establishing some simple models of local cortical dynamics (14). Less is known about the impact of this relationship for downstream circuits (though see (15)).

As an example, consider the detection of a sensory stimulus, which has been foundational in the human (16-21) and non-human primate psychophysical literature (22,23) and serves as one of the most widely utilized behavioral paradigms in rodent literature (24-26). In an attempt to link the underlying neural variability to behavior, the principal framework for describing sensory perception of stimuli near the physical limits of detectability is signal detection theory (27). A key prediction of signal detection theory is that, on single trials, detection of the stimulus is determined by whether the neural response to the stimulus crosses a threshold. Particularly large responses would be detected but smaller responses would not, so variability in neural responses would lead to, and perhaps predict, variability in the behavioral response. From the perspective of an ideal observer, if variability in the sensory-evoked response can be forecasted using knowledge of cortical state, the observer could potentially make better inferences, but in traditional (state-blind) observer analysis, the readout of the ideal observer is not tied to the ongoing cortical state.

In this work, using network activity recordings from the whisker sensitive region of the primary somatosensory cortex in the awake mouse, we develop a data-driven framework that predicts the trial-by-trial variability in sensory-evoked responses in cortex by classifying ongoing activity into discrete states that are associated with particular patterns of response. The classifier takes as inputs features of network activity that are known to be predictive of single-trial response from previous studies (9,14), as well as more complex spatial combinations of such features across cortical layers, to generate ongoing discrete classifications of cortical state. We optimize the performance of this state classifier by systematically varying the selection of predictors. Finally, embedding this classification of state in a state-aware ideal observer analysis of the detectability of the sensory-evoked responses, we analyze a downstream readout that changes its detection criterion as a function of the current state. We find that state-aware observers outperform state-blind observers and, further, that they equalize the detection accuracy across states. Downstream networks in the brain could use such an adaptive strategy to support robust sensory detection despite ongoing fluctuations in sensory responsiveness during changes in brain state.

## Results

To directly assess the relationship between ongoing cortical activity and variability in the sensory-evoked cortical response, we recorded extracellular activity across layers of cortex in the awake head-fixed mouse. Specifically, spontaneous and sensory-evoked local field potentials (LFPs) were recorded using a 32-channel laminar array targeted to the region of the primary somatosensory cortex corresponding to facial vibrissae (S1 barrel cortex, Fig. 1A). Mice were subjected to brief single-whisker deflections (11 recordings in 6 mice; average 438 (196 – 616) trials per recording). Intrinsic optical signal imaging was performed to locate the barrel column corresponding to the stimulated whisker. The sensory stimulus (Fig. 1B, top) was a computer-controlled punctate deflection in the caudo-rostral plane (see Methods), designed to emulate velocity transients observed during ‘stick-slip’ events in rodents whisking across surfaces (28–30). Cortical layers were assigned based on the trial-averaged spatial profile of the sensory-evoked LFP and current source density (CSD) responses, with layer 4 centered on the largest evoked response in the LFP and the large, early current sink in the CSD (Figure 1C, see Methods). Putative boundaries between layer 4 and layer 2/3 or layer 5 were based on published laminar dimensions (31).

**Figure 1.**
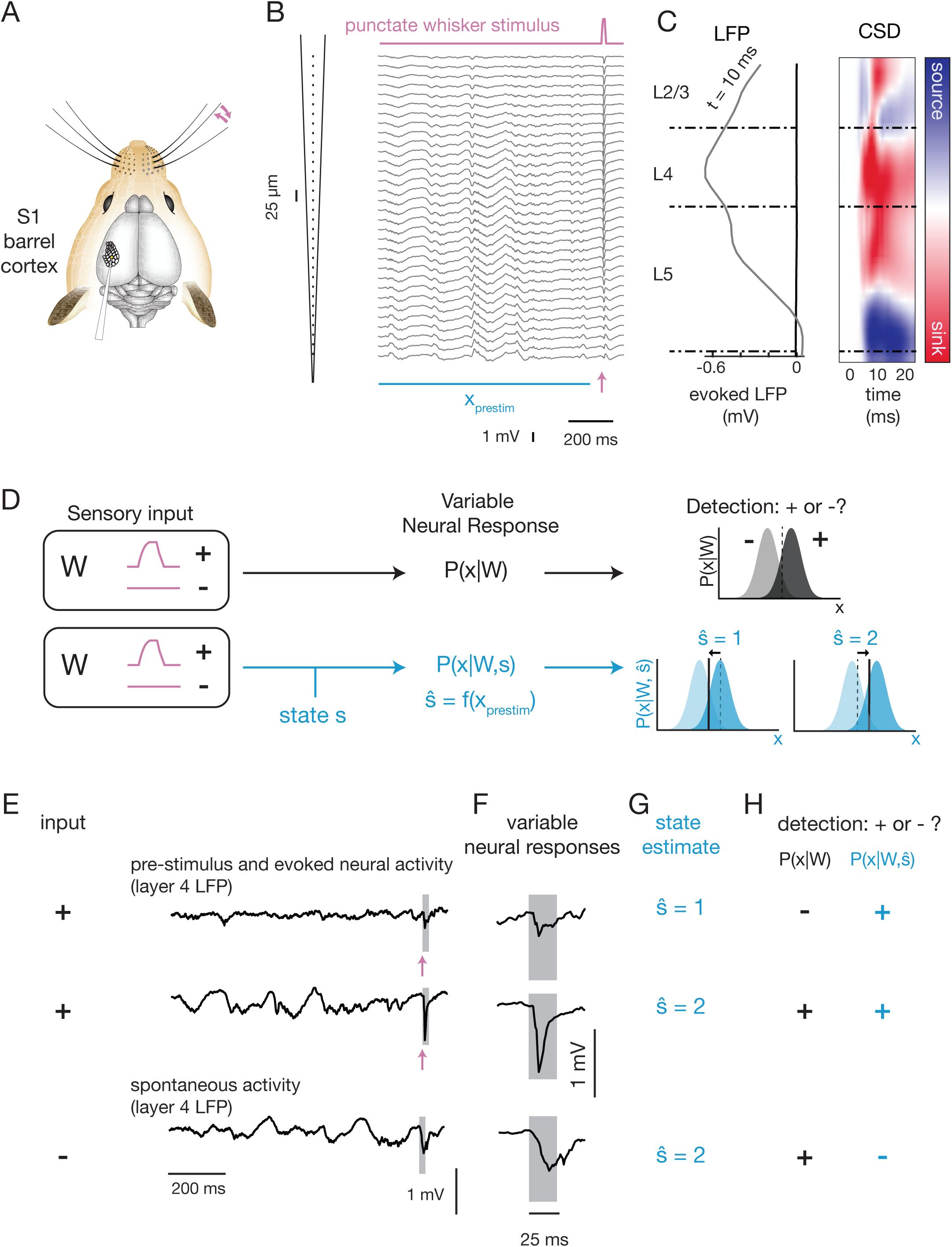
Single-trial prediction and detection of sensory-evoked responses in the awake mouse. **A.** Recordings of whisker-evoked neural activity in the principal barrel column were acquired using a laminar probe in awake mice. **B.** A single trial response to the punctate whisker stimulus (galvanometer trace, pink line at top), showing LFP activity across the 32 channels of the silicon probe (25 um spacing), with 1 s of prestimulus activity (*x*_*prestim*_, horizontal blue bar) and the sensory-evoked response starting at the time point indicated by the arrow (pink). **C.** Average spatial profile of evoked LFP at t = 10 ms post-stimulus (left). Right: average evoked CSD over the 25-ms post-stimulus window, with color indicating whether sink (red) or source (blue). Channels were assigned to layers (dashed horizontal lines separating sections labeled L2/3, L4 and L5) by centering the largest evoked response in layer 4. **D.** State-blind analysis (top row, black font): Neural responses encode sensory inputs, described by the encoding distribution P(x|W) of response (x) conditioned on stimulus identity (W, here the presence (+) or absence (-) of a sensory input). An ideal observer determines the stimulus from the response using P(W|x). State-aware analysis (bottom row, blue font): sensory responses also depend on cortical state (s). Here we estimate ŝ by computing a function of the pre-stimulus neural activity (x at time t’ < t). An ideal observer sets a state-dependent threshold depending on the estimated state the estimated state. Fig. 2-4: estimation of ŝ. Fig. 5-6: state-aware ideal observer analysis using P(W|x, ŝ) **E-H.** Illustration of the detection framework applied to an example drawn from recorded data. **E.** The recorded LFP in layer 4 is shown for three example trials, with W = + (top two) and W = (bottom). **F.** Expanded view of neural activity in a 25-ms window (gray boxes in (E)). In these selected examples, spontaneous fluctuations (W = -) are larger than the first sensory-evoked response (W = +). **G.** The state classifier (described in Fig. 2-4) estimated state ŝ for each trial based on ongoing/prestimulus activity. **H.** Illustration of detection of input (W = + or W = -) based on state-blind threshold choice (black, P(x|W)) and based on state-aware threshold choice (blue, P(x |W, ŝ)). In this example, the threshold for state 2 was increased to reject the spontaneous activity (bottom row), while the threshold for state 1 was decreased to accept the small sensory-evoked responses.

Sensory-evoked responses in layer 4 were variable: mean amplitude of the response in layer 4 was a negative dip of 0.81 mV (+/− 0.35 across recordings, N = 11), and the standard deviation of evoked response size across trials was 0.45 mV (average SE across recordings, N = 11). We examined the impact of such variability on the detectability of sensory inputs in the framework of ideal observer analysis, which is conceptually presented in Figure 1D. In this scenario (Fig. 1D, top row), a sensory stimulus *W* takes on one of two possible values: “+” in the case that a sensory input was present and “−” in the absence of a sensory input. Neural activity (x, see Methods), either spontaneous (W is “−”) or generated by the stimulus (W is “+”), was variable across trials and described by a conditional distribution *P*(*x|W*). The task of a downstream network, imagined here as an ideal observer, was to determine from the neural activity whether or not there was a stimulus. In the classical signal detection framework, this was envisioned as the observed activity arising from one of two distributions, *P*(*x*|*W* = “−”) or *P*(*x|W = “+”)*. The ideal observer ascribes activity above a chosen threshold (to the right of dashed red line) as belonging to *P*(*x|W = “+”)*, and thus concludes that a stimulus was present, and otherwise as belonging to *P*(*x|W = “−”)*, and thus the stimulus was deemed absent. In this work, we considered an alternative perspective, which is that the ideal observer was “state-aware”. That is, we considered the case in which the response distribution *P*(*x|W, s)* depended upon ongoing activity (“state,” s) as well as the stimulus (Fig. 1D, blue, bottom row). In this case, the discrete state (*ŝ*) is classified from the recorded, ongoing cortical activity, which is subsequently used by the state-aware observer to set the detection threshold independently for each state.

To illustrate how this framework operates, we show a set of example trials from a single recording in Fig. 1E-H. Two of these examples of layer 4 LFP activity show responses to a whisker input and one is a segment of spontaneous activity (Fig. 1E). Across the stimulus-evoked responses, we observed significant variability in the overall size of sensory-evoked response (Fig. 1F). Moreover, one of the evoked responses (Fig. 1F, top row) is smaller than a spontaneously occurring LFP event (Fig. 1F, bottom row). We also note that pre-stimulus LFP activity is quite different across these recordings. The goal of the state classifier is to find consistent relationships between features of the ongoing activity and the details of the single-trial sensory-evoked response. Assuming for the moment that it is able to do so, the state classifier would classify the pre-stimulus activity for these responses (Fig. 1E) into separate states (*ŝ* = 1 or 2, Fig. 1G). When tasked with detecting or rejecting stimuli on the basis of the LFP response (Fig. 1H), an ideal observer sets a single threshold for detection, which causes it to fail to detect a true sensory response while generating a false alarm on the spontaneous fluctuation (Fig. 1H, black). In contrast, the state-aware observer sets its detection criterion separately for each state. For this example, the threshold may be lowered for state 1 and raised for state 2, thus the sensory-evoked responses would be detected but the spontaneous fluctuation would be rejected (Fig. 1H, blue).

The state-aware observer thus has two distinct stages: state classification and sensory stimulus detection. The idea is that, by adapting its criterion for detection in accordance with the expected response, the state-aware observer will more reliably detect sensory-evoked responses and reject spontaneous fluctuations. Overall, the success of this strategy depends on a state classifier that predicts variation in the future sensory-evoked response, so we first optimized classification models with the goal of identifying the most relevant features of ongoing activity for the prediction of the details of the sensory-evoked responses. We then use this framework to classify ongoing activity into states and compare traditional (state-blind) and state-aware observers to determine how using this prediction to adjust detection strategy impacts overall detection performance.

### State classification based on prediction of cortical responses

The foundation upon which the state-aware observer is constructed is a prediction of the sensory-evoked cortical response. This prediction is based on classifying elements of the ongoing, pre-stimulus activity into discrete “states,” and the goal is to find the features of ongoing activity and the classification rules that generate the best prediction of sensory-evoked responses. The features of ongoing activity include the power spectrum of pre-stimulus LFP and the instantaneous “LFP activation” (Fig. 2A). To describe sensory-evoked responses, we define a parameterization of the LFP response using principal components analysis (Fig. 2B). The state classifier is a function that takes as inputs features of pre-stimulus LFP and produces an estimate of the principal component (PC) weights and thus of the single-trial evoked response (Fig. 2C). In the following sections, we describe this process in detail.

**Figure 2.**
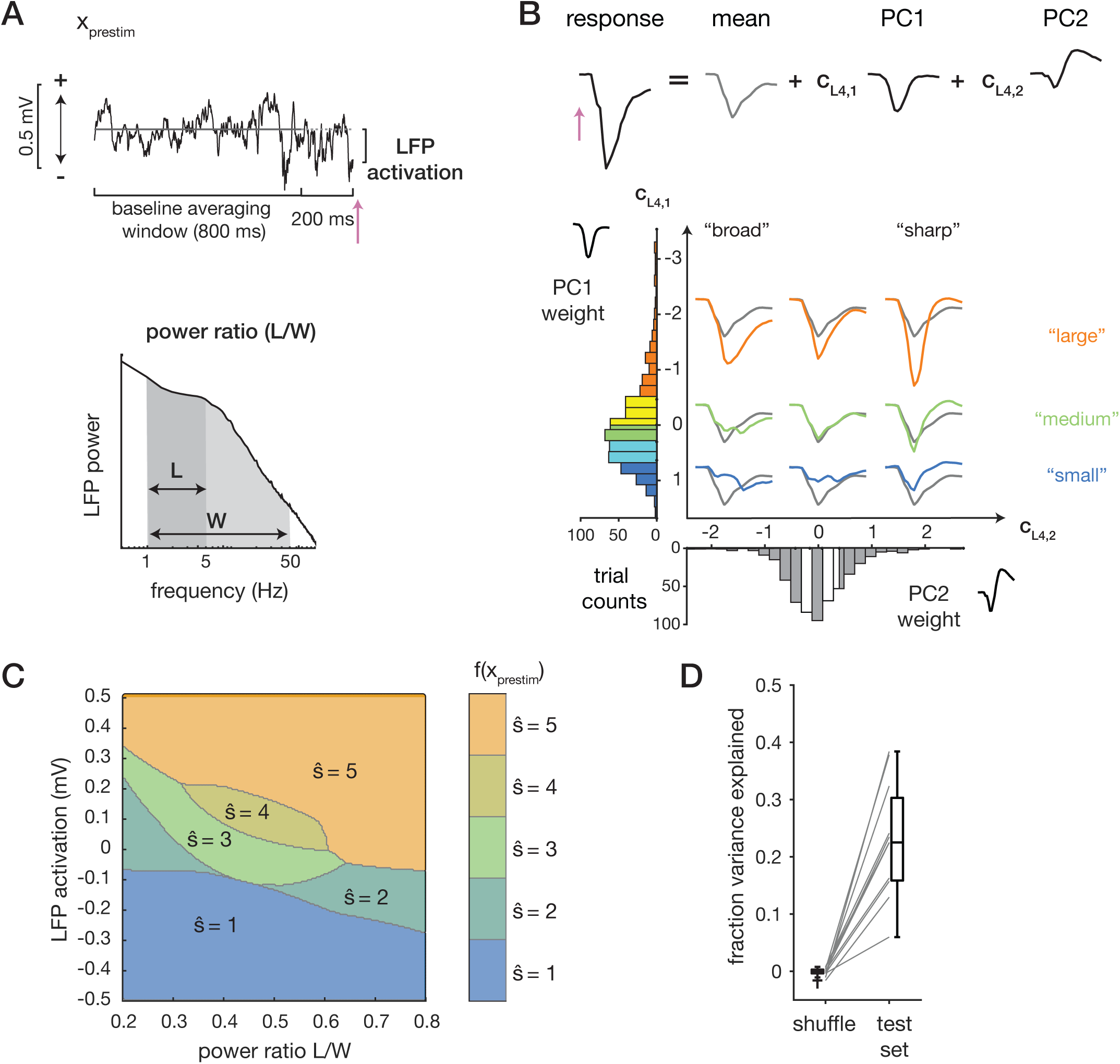
A state classifier to predict single-trial sensory-evoked LFP responses. **A.** Features of the pre-stimulus activity used for the prediction. Top row: LFP activation is the LFP in the 10 ms preceding stimulus onset (pink arrow) minus the LFP averaged over the baseline window. Bottom row: power ratio is the ratio of LFP power in the 1-5 Hz range (L) to the power in the 1-50 Hz range (W). **B.** Top row: parameterization of the early (25-ms) sensory-evoked LFP response through principal components. Bottom row: Illustration of evoked responses represented in PC1-PC2 weight space. Five ranges are defined based on quintiles. Color indicates the PC1 ranges: largest responses, orange; smallest responses, blue. Averages of responses falling into the 1st, 3rd, and 5th quintiles of PC1 weights and PC2 weights are plotted, colored by the PC1 quintile (orange, green, blue) with the overall average evoked response in gray. **C.** Classifier boundaries in power ratio-activation space that predict the PC1 weights. Regions are labeled from 1 (“small”) to 5 (“large”) and colored as in panel B. See 5upporting Figures 1 and 2 for boundaries for other recordings. **D.** Fraction of variance explained by the classifiers on trial-shuffled data and on the test set.

### Initial selection of features of ongoing activity

The first step was to select features of ongoing activity, which are the inputs to the state classifier. Based on previous work in somatosensory cortex (8,9), we first identified two features that reflect the instantaneous phase and synchronization of ongoing activity measured through LFP. To capture instantaneous phase, we used pre-stimulus LFP activation, defined as the average LFP recorded over the 10 ms preceding the stimulus, relative to the baseline activation level (LFP averaged from -1000 ms to -200 ms, Fig. 2A, top). Synchronization is reflected by the low-frequency power in the LFP, quantified as the ratio of power between 1 and 5 Hz (denoted here as “L”) to the total power between 1 and 50 Hz (denoted here as “W”, for wideband, Fig. 2A, bottom). Power ratio was computed using the fast Fourier transform over the 2-second pre-stimulus period.

### Parametrizing variability in evoked responses

The next step was to define a discrete parameterization of the sensory-evoked LFP response, which will be the targeted output of the state classifier. We focused on the single-trial sensory evoked LFP response in cortical layer 4 in a 25-ms period following the delivery of the sensory stimulus. We parameterized the evoked response using the principal components (PCs) of the evoked response across trials. On average, the first two modes accounted for 89 % ± 2 % (N=11 recordings) of the cross-trial variance. Each trial was therefore described as the average response, plus the weighted sum of the first two PCs (Fig. 2B, top, Methods). Variation in the weight of PC1 (*c*_*L*4,1_) captured differences in the overall amplitude of the response, while variation in the weight of PC2 (*c*_*L*4,2_) captured the temporal width of the response (Fig. 2B). Thus, the PC representation captured recognizable characteristics of the shape and size of the evoked response.

To discretize the parameterization, we defined five evoked response ranges for each PC weight based on quintiles of their distributions (Fig. 2B), and set the weight for any trial falling within a given range to the average of the individual weights within the range (i.e. the quintile average; see Methods). To quantify how much discretization impacts prediction power, we computed the variance explained by the discrete and continuous parameterization. The discrete parameterization captures an average of 78 % ± 4 % of variance, compared to 89 % ± 2 % (N = 11 recordings) for the continuous representation. This number sets the maximum fraction of variance explained that could be achieved by any state classifier that generates a prediction of the evoked response.

### Building the state classifier

We fit classifiers that mapped combinations of pre-stimulus features to specific ranges of the parameterized responses (Figure 2C). This determines a set of boundaries that define the pre-stimulus states. For the example recording shown in Fig. 2C, smaller evoked responses were associated with negative pre-stimulus LFP activation (Fig. 2C, blue, *ŝ*_1_ = 1), while large evoked responses were associated with positive pre-stimulus LFP activation (Fig. 2C, orange, *ŝ*_1_ = 5). Boundaries found for states were similar across different recordings (Supplemental Figure 1-2). This finding is consistent with previous reports of the negative interaction between ongoing and evoked activity (32): sensory evoked responses were smaller when the ongoing activity was higher.

Prediction performance was quantified using the fraction of variance explained (fVE) over the 25-ms evoked response window (Methods). We partitioned the data into three sets: one for fitting classifiers (“training”), one for selecting the best pre-stimulus features (“cross-validation”) and one for quantifying fVE (“test”). This was important to avoid spuriously high performance of optimized classifiers due to over-fitting (see Methods for details). Using activation and power ratio (as defined in Fig. 2A) as pre-stimulus features, we found that the fVE was 0.18 ± 0.03, which was significantly larger than the fVE when trials were shuffled within each recording (average fVE for trial-shuffle: 0.001 ± 0.002, N = 11 recordings; Fig. 2D).

### Optimization of prediction of the single-trial sensory-evoked response

Next, within the general class of pre-stimulus features considered power ratio and LFP activation we optimized several choices: the range of frequencies used to compute the power ratio; the cortical depth from which the ongoing LFP signal is taken; and possible combinations of LFP signals across the cortical depth. Changes in pre-stimulus features resulted in changes in the boundaries between states, and ultimately in changes in prediction performance.

First, we varied the bounds of the low-frequency range (“L range”, Fig. 3A). The increase in fVE was on average 0.09 ± 0.05 (N = 11 recordings) (Fig. 3B; classifier boundaries shown in Supplemental Figure 2), with a significant increase in 10 of 11 recordings (Fig. 3C, asterisks). We found that the optimal L range could extend to frequencies up to 40 Hz (Fig. 3C), with the median bounds of the optimal L being from 1 to 27 Hz.

**Figure 3.**
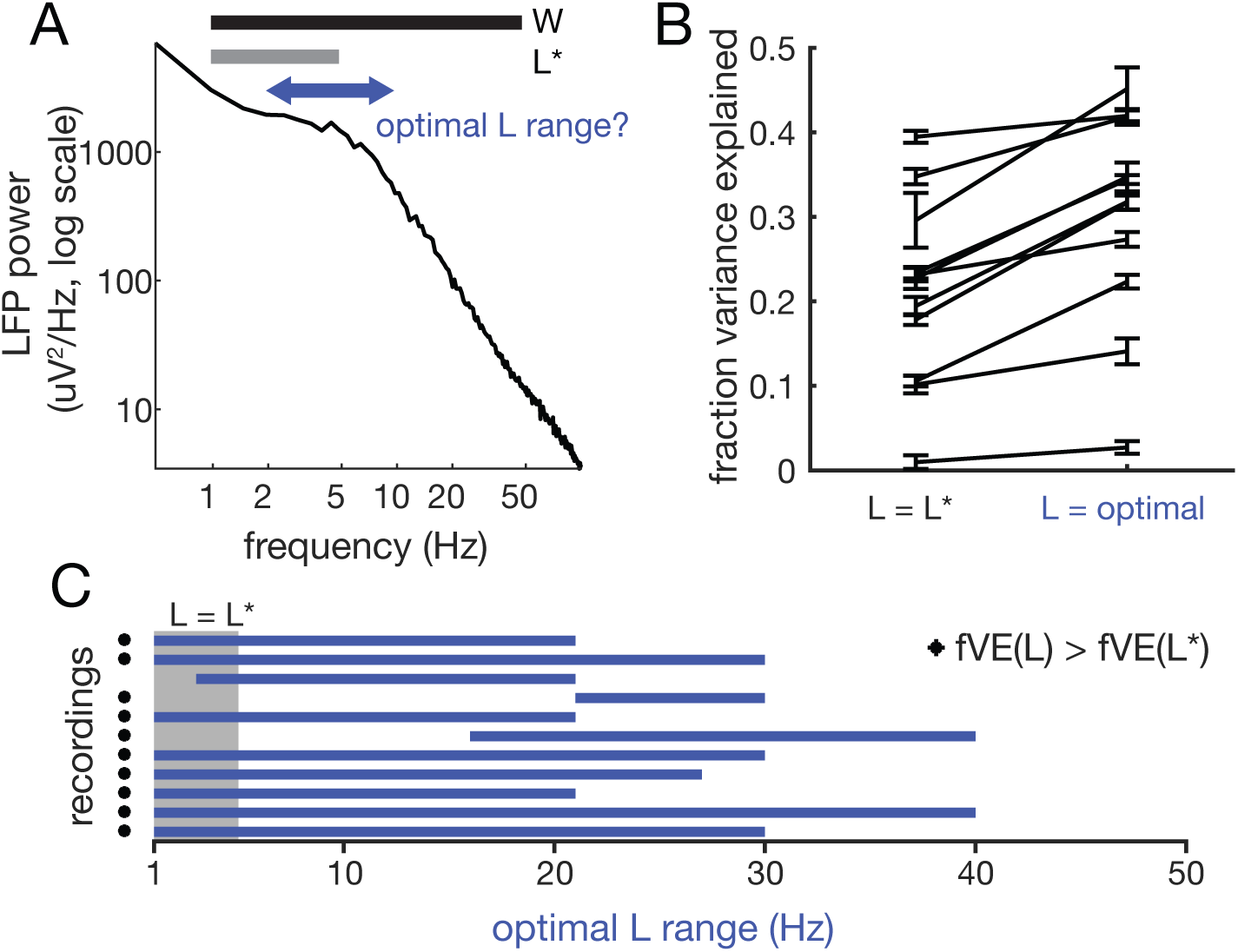
Optimizing state classifier performance: defining the power ratio. **A.** Optimization of power ratio windows. We optimized the classifier by varying the ranges of low-frequency power used to compute power ratio, while keeping W fixed. **B.** Fraction of variance explained by the state classifier based on original power ratio (L*/W) and optimal (L/W). Error bars (SE) are bootstrap estimates of uncertainty in fVE calculation. **C.** Optimal L range across all recordings (blue bars). Gray box shows the original range for L (1-5 Hz). Asterisks indicate those recordings for which the increase in fVE was > 2 SE.

Using for each recording the power ratio based on the optimized range of low-frequency power (Fig. 3), we next determined where along the cortical depth the most predictive activity was and whether taking spatial combinations of LFP activity could improve the prediction. Note that in this analysis, the channel for the stimulus-evoked response was held fixed (L4) and thus the parameterization of the evoked response using principal components did not change, but the pre-stimulus channel was varied. For each recording, we thus built a series of classifiers, using single- and multi-channel LFP activity from across the array (Fig. 4A, Supplemental Figure 3), which again were optimized for prediction of the single-trial L4 sensory-evoked response.

**Figure 4.**
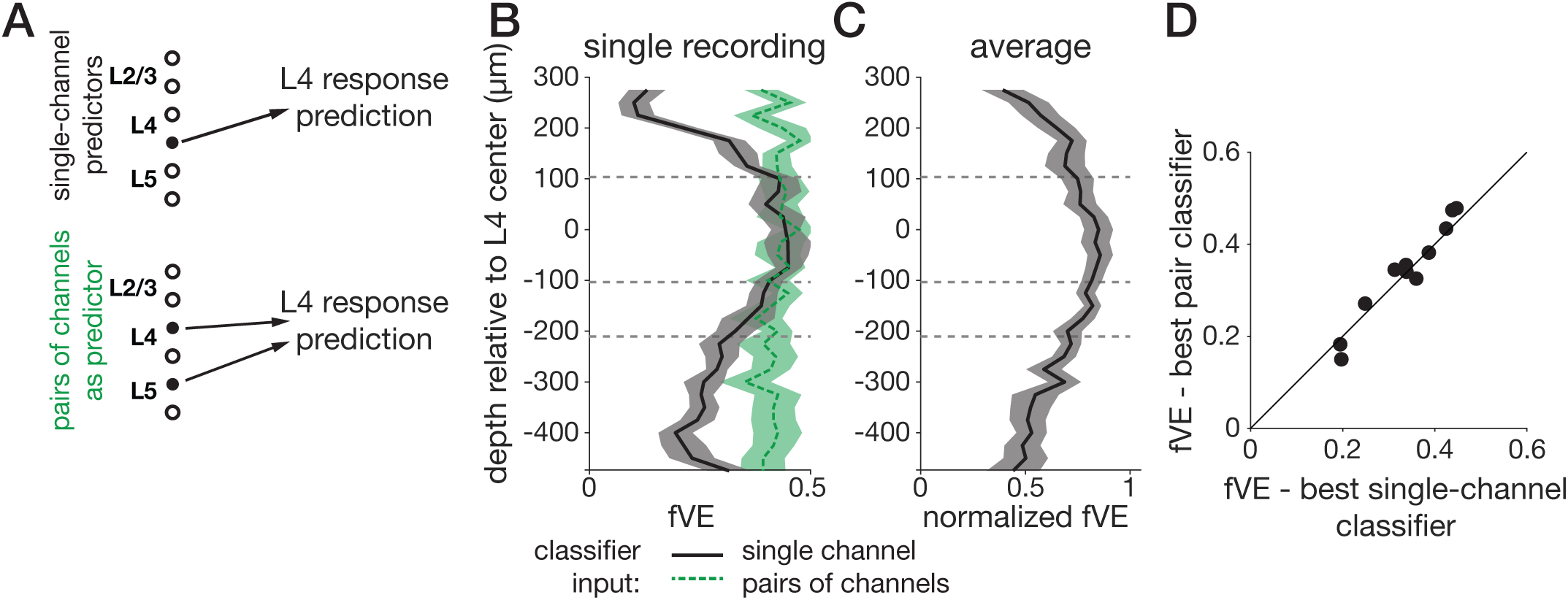
LFP recorded near layer 4 drives optimal state classifier performance. **A.** We constructed two sets of state classifiers: first, using the power ratio and activation from single channels across the array (black label, top) and second, by adding to the best single-channel classifier a third predictor, the activation from a secondary channel (green label, bottom). **B.** Spatial profile of fVE for a single recording. Each classifier took as input one of the recorded channels. Vertical axis shows the depth of the input channel with fVE on the horizontal axis. Shaded regions show ± 1 SEM estimated from bootstrapped fVE calculation. Black, solid: single channel LFP. Green, dash: classifiers based on pairs of channels. **C.** Normalized fVE spatial profile averaged across recordings, with fVE normalized by the maximum fVE on each single recording. Recordings were aligned based on assigned L4 channels and then averaged. Each point is an average over N = 9 recordings with > 500 µm of overlap after alignment. Shaded region shows ± 1 SEM across recordings. **D.** Performance of classifiers using pairs of channels vs. the best single-channel classifier for N=11 recordings. Estimated error bars (from bootstrap) are approximately ± 0.02 and omitted for clarity. No significant difference between single-channel vs. pair-channel classifiers for any recording.

Classifiers built from a single channel of LFP performed best when the channel was near L4 (Fig. 4B, single example; 4C, average profile). Because the LFP represents a volume-conducted signal, we also examined the current source density (CSD) (33-35), estimated on single trials using the kernel method (Potworowski et al. 2012). There was no improvement in fVE using CSD to build classifiers (fVE difference, CSD minus LFP: -0.07; range: (−0.12, -0.01)).

For each recording, we defined an optimal classifier channel based on the spatial profile of fVE for single-channel predictors (Fig. 4B; Supplemental Figure 3). In the “pair” combination, we paired the optimal classifier channel with each of the other possible 31 channels (Fig. 4B; green dashed line). We optimized the classifier in the 3-dimensional space defined by power ratio (on the optimal channel only) and LFP activation from each of the two channels and compared the fVE to that obtained using the optimal classifier channel only (Fig. 4D). We found no improvement in the prediction using the pair combination compared to using the optimal channel alone (Fig. 4D, mean fVE difference: 0.00 ± 0.01; 0/11 recordings with significant change, pair vs. single) or using more complex combinations of channels (Supplemental Figure 3).

To summarize, we optimized classifiers based on pre-stimulus features to predict single-trial sensory-evoked LFP responses in S1 cortex of awake mice. We found that the classifier performance was improved by changing the definition of the power ratio (L/W) such that the low-frequency range (L) extended from 1 Hz to 27 Hz, depending on the recording, which differed from the range typically used from anesthetized recordings in S1 (1-5 Hz) (8,9). We also found that the most predictive pre-stimulus LFP activation was near was near layer 4.

### Ideal observer analysis of sensory-evoked responses

After establishing a clear enhanced prediction of the single-trial stimulus-evoked response within the LFP by considering the pre-stimulus activity, we investigated the impact of this relationship on the detection of sensory stimuli from cortical LFP activity using a state-aware ideal observer analysis. We first considered a simple matched-filter detection scheme (37) in which the ideal observer operated by comparing single-trial evoked responses to the typical shape of the sensory evoked response (Methods, *Detection*). The matched filter was defined by the trial-average evoked LFP response, and this filtered the raw LFP (Fig. 5A) to generate the LFP score (Fig. 5B). For the state-blind observer, a detected event was defined as a peak in the LFP score that exceeded a fixed threshold (Fig 5B, stars). The LFP score distributions from time periods occurring during known stimulus-evoked responses and from the full spontaneous trace were clearly distinct but overlapping (Fig. 5C), and detected events (Fig. 5B, stars) included both “hits” (detection of a true sensory input) and “false alarms” (detection of a spontaneous fluctuation as a sensory input).

**Figure 5.**
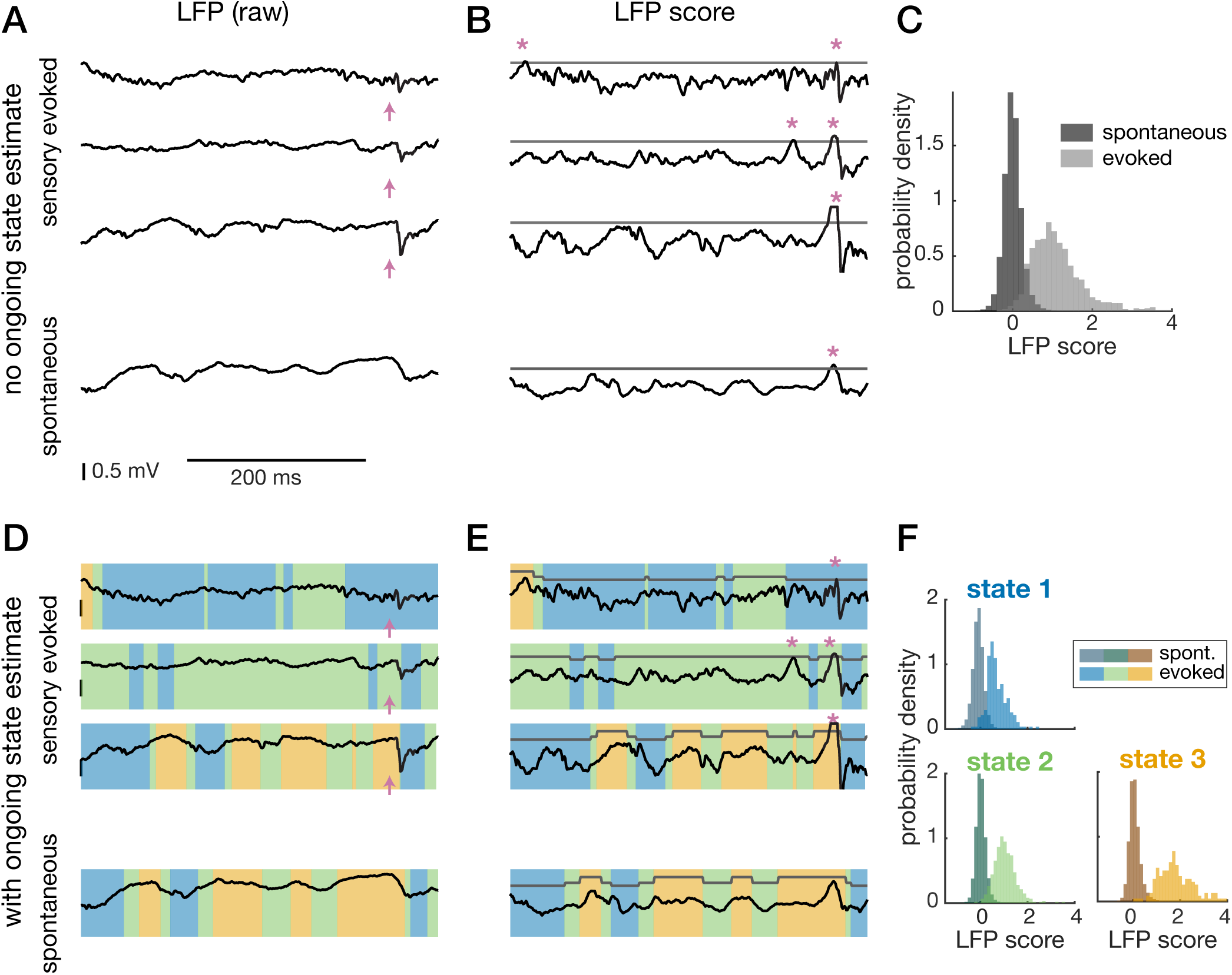
State-aware ideal observer analysis of spontaneous and sensory-evoked activity. **A.** Example of a 400-ms epoch of LFP taken from sensory-evoked trials (top 3) and spontaneous periods (bottom). Pink arrows indicate stimulus delivery time. **B.** LFP traces convolved with a filter matched to the average evoked response generate the LFP score. Peaks in the score that are above a threshold (gray) are detected as events (pink asterisks). **C.** Distribution of scores during spontaneous activity (dark gray) and of sensory-evoked responses (light gray). The two distributions are clearly distinct but with an overlapping area, leading to spontaneous events being detected as sensory-evoked responses (false alarms). **D.** Same example traces as in A, but with the “state” at each point in time indicated by the color overlay. **E.** An example of how the detection threshold could be adjusted depending on the value of the ongoing state. **F.** Separating by ongoing state generates three sets of spontaneous/evoked score distributions (analogous to C).

Next, using the state classifier constructed in the first half of the paper, we analyzed the performance of a state-aware observer on a reserved set of trials, separate from those used for fitting and optimizing the state classifiers (Methods). Specifically, using the optimized state classifier (Figs 3 and 4), we continuously classified “state” at each time point in the recording (Fig. 5D). The state-aware observer detects events exceeding a threshold, which changed as a function of the current state (Fig. 5E). Instead of a single LFP score distribution, we now have one for each predicted state (Fig. 5F), leading to many possible strategies for setting the thresholds for detecting events across states.

In general, the overall hit rate and false alarm rate will depend on hits and false alarms in each individual state (Fig. 6A-B), as well as the overall fraction of time spent in each state (Fig. 6A, inset). To compare between traditional (state-blind) and state-aware observers, we compared hit rates at a single false alarm rate, determined for each recording as the false alarm rate at which 80%-90% detection was achieved by a state-blind ideal observer. To select thresholds for the state-aware observer, we systematically varied the thresholds in state 1 and state 3, while adjusting the state-2 threshold such that average false alarm rate was held constant. For each combination of thresholds, we computed the overall hit rate (Fig. 6C). For this recording, the state-aware observer (hit rate: 96%) outperformed the traditional one (hit rate: 90%). This worked because the threshold in state 3 could be increased with very little decrease in the hit rate (Fig. 6B), and this substantially decreased the false alarm rate in state 3 (Fig. 6A). Because the *overall* false alarm rate is fixed, this meant more false alarms could be tolerated in states 1 and 2. Consequently, thresholds in states 1 and 2 could be decreased, which increased their hit rates. Across recordings, we found that the state-aware observer outperformed the state-blind observer in 9 of 11 recordings (Fig. 6D; Supplemental Figure 4-5). Hit rates slightly but significantly increased from a baseline of 81% for the state-blind observer to 84% for state-aware detection, or an average change of +3 percentage points (SE: 3%; signed-rank test, p < 0.01, N = 11). Although this is a modest change, overall hit rates were computed from the average of hit rates in each state, weighted by the amount of time spent in each state. To separate these factors, we analyzed the hit rate of the state-blind and state-aware observers by computing, for each observer, the hit rate conditioned on each pre-stimulus state (Fig 6E). For this recording, the state-blind observer had very low hit rate in state 1 and high hit rates in states 2 and 3. In comparison, hit rates were similar across the three state for the state-aware observer (Fig. 6D). Thus, in state 1 (smallest responses, blue), we observed a large increase in the hit rate depending on whether the observer used state-blind or state-aware thresholds. Averaged across all recordings, the state-1 hit rates increased from 60% to 76%. This is a relative increase of 26% (SE 11%), which is substantial. Because this is weighted by the fraction of time spent in state 1, the overall impact on the hit rate is smaller. Hit rates increased slightly on average in state 2 (+ 2%, SE 4%) and decreased slightly in state 3 (−7%, SE 9%). The net impact of this is that across the majority of recordings, the cross-state range of hit rates for the state-blind ideal observer was much larger than that for the state-aware ideal observer (Fig. 6D, Fig. 6F; 19%, average state-blind minus state-aware hit rate range in percentage points (SE: 5%); p < 0.01, signed-rank test, N = 11). Thus, the state-aware observer has more consistent performance across all pre-stimulus states than a state-blind observer.

**Figure 6.**
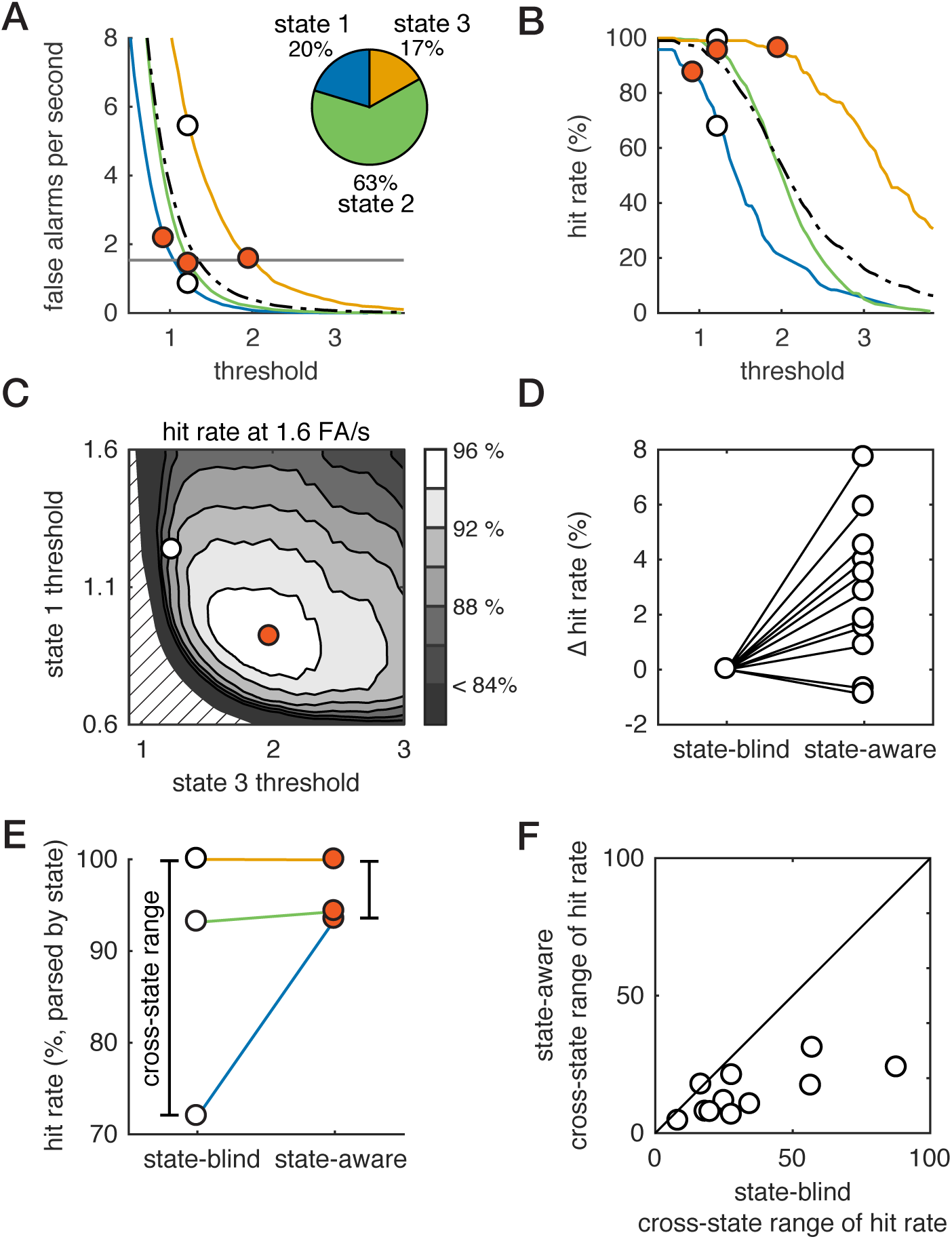
Optimizing thresholds for the state-aware observer. **A.** False alarm (FA) rate vs. threshold for each pre-stimulus state (colors) as well as the full dataset (black dashed). Pie chart: fraction of time spent in each state. White fill circles: threshold for state-blind observer. Vermillion fill circle: threshold for state-aware observer (see C). Combined with the fraction of time spent in each state (pie, A), this sets the overall FA rate. **B.** Hit rate vs. threshold for each pre-stimulus state (colors) as well as the full dataset (black dashed). Combined with the fraction of time spent in each state (pie, A), this sets the overall hit rate. Circles marks same thresholds as in (A). **C.** Hit rate of state-aware observer for combinations of threshold that generate the same (1.6 Hz) FA rate is indicated by grayscale. Circles correspond to the combinations of thresholds shown in A and B. Dashed lines show region with no solution satisfying constraint on FA rate. **D.** Change in hit rate between state-blind and state-aware observers. Thresholds are chosen based on the optimization shown in (C), and hit rates are computed on a reserved set of trials and compared to the fixed-threshold hit rate at equivalent false alarm rate. **E.** For the example recording, we parse hit rate by pre-stimulus state for the state-blind and state-aware observers. The cross-state range of hit rates is defined as the range of hit rates observed across the three states. **F.** Cross-state range of hit rates for the state-aware observer versus cross-state range for the state-blind observer (see D). The cross-state range is smaller for the state-aware observer, meaning that hit rates are more consistent across pre-stimulus states for the optimal state-aware observer.

## Discussion

Due to the rapid development of tools that enable increasingly precise electrophysiology in the awake animal, there is a growing appreciation that the “awake brain state” encompasses a wide range of different states of ongoing cortical activity, and that this has a large potential impact on sensory representations during behavior (38-43). Here, we constructed a framework for the prediction of highly variable, single-trial sensory-evoked responses in the awake mouse based on a data-driven classification of state from ongoing cortical activity. In related work, past studies have used some combination of LFP/MUA features to successfully enhance the prediction of future evoked MUA response (9,14). In this study, we extended these types of approaches for state classification and response prediction to cortical recordings in the awake animal, opening up the problem to allow complex combinations of ongoing activity in space as well as different features of the power spectrum of the pre-stimulus activity. We found that simple features of pre-stimulus activity sufficed to enable state classification that yielded single-trial prediction of sensory evoked responses. These predictive features were analogous to the synchronization and phase variables found in previous studies (8,9,14), though we found a revised definition of synchronization was more predictive. In particular, we found that the very low-frequency band of the LFP power spectrum (1-5 Hz) was less predictive of single-trial evoked responses in our recordings than a wider band (e.g. 1 to 27 Hz). This is consistent with findings from a recent study (39) that surveyed the power spectrum of LFP across different behavioral states in the awake animal and demonstrated differences in the power spectrum between quiet and active wakefulness up to 20 Hz. While we have focused on the problem of state classification and prediction from the perspective of an internal observer utilizing neural activity alone, future work could investigate whether the state classifier is also tracking external markers of changes in state, such as those indicated by changes in pupil diameter (41,44), whisking (39), or other behavioral markers in the awake animal.

In its current formulation, this framework utilizes only the features of ongoing cortical activity that are reflected in the LFP in order to classify state and predict the evoked LFP response. Both as features underlying the state classifier and as the sensory-evoked response being predicted, LFP must be interpreted carefully, as the details of how underlying sinks and sources combine depend on the local anatomy and population spiking responses (45). In barrel cortex, the early whisker-evoked LFP response (0 to 25 ms) is characterized by a current sink in L4 initially driven by thalamic inputs to cortex, but also reflecting cortical sources of activity: the evoked LFP is highly correlated with the layer-4 multi-unit activity response (46,47). We restricted our predictive framework to the high degree of variability in this initial response. It remains to determine how LFP response variability is reflected in the sensory-evoked single-unit cortical spiking activity patterns. Further, regarding LFP as a predictor used by the state classifier, LFP is a mesoscopic marker of cortical state that neglects finer details of cortical state organization. In addition to establishing whether better predictions are made from more detailed representations of cortical state, it is an interesting question how microcircuit spiking dynamics are related to the mesoscopic markers of cortical state, or how much can be inferred about population spiking dynamics from the LFP. Finally, thalamic and cortical activity are tightly linked, and the results presented here may also reflect variations in ongoing thalamic activity. Disentangling thalamic and cortical sources of variability in the evoked response will require paired recordings and perturbative experimental approaches designed to address issues of causality.

In the second part of the paper, we used ideal observer analysis to show that state-aware observers, with oracle knowledge of the spontaneous, ongoing state fluctuations informative of the single-trial sensory-evoked response, can out-perform a state-blind ideal observer. Our analysis relied on classification of the markers of ongoing state. This is not to suggest that this specific estimation takes place in the brain, but instead could potentially be achieved dynamically by a downstream network through the biophysical properties of the circuitry. Theoretically, the gain and threshold of such a readout neuron or network could be dynamically modified on the basis of the ongoing activity as a biophysical manifestation of the adaptive state-aware ideal observer, though the identification of specific mechanisms was beyond the scope of the current study.

We found that the state-aware observer had higher accuracy than the traditional, state-blind observer, but the absolute gain in hit rate (at fixed false alarm rate) averaged across all states was modest. When pre-stimulus states were analyzed separately, however, we found that accuracy in the low-response state was substantially higher for the state-aware observer, where there was a relative increase of 25% in the hit rate for this state. Because small sensory responses are predictable from the ongoing activity, transiently lowering the threshold for detection resulted in more “hits” in the low-response state, while false alarms in high-response states could be avoided by raising the threshold when the state changed. However, the cortical activity was classified to be in this particular state only approximately 20% of the time, and thus had a relatively modest impact on the overall performance, averaged across all states. What is not currently known is the overall statistics associated with the state transitions (i.e. distribution of time spent in each state, rate of transitions, etc.) during engagement within perceptual tasks, but in any case, what we observe here is a normalization of detectability across brain states.

For near-threshold sensory perception, the signal detection theory framework asserts that single-trial responses are predictive of perceptual report (27). While there are many previous studies that seem to support this (48-51), several animal studies have called this into question, showing that primary sensory neural activity does not necessarily co-vary with perceptual report on simple detection tasks (22,24,26). It is entirely possible that the conflicting findings in the literature are due to behavioral state effects, and that more consistent reports would emerge if the analysis of the neural activity incorporated elements of the state-classification approach developed here. Our results show how single-trial response size can be decoupled from perception, if a downstream network predicts and then accounts for the variability in sensory responses. Moreover, our analysis showed that some states of pre-stimulus activity should be associated with higher or lower performance on a near-threshold detection task, which has been observed in near-threshold detection studies in the rodent (25) and monkey (23). It should be noted that there is controversy regarding the relevance of primary sensory cortex in simple behavioral tasks (52,53), but this is likely related to the task difficulty (54), where a large body literature has resolutely shown that processing in primary cortical areas is critical for difficult tasks that increase cognitive load, and we suspect that near threshold stimuli such as those shown here fall in that category.

Many studies have demonstrated a link between pre-stimulus cortical activity and perceptual report on near-threshold detection tasks in humans (16,17,55-58). Currently, it is not entirely clear how far the parallel in cortical dynamics between the mouse and human can be taken. One challenge is that connecting invasive recordings in the mouse to non-invasive recordings in human studies is non-trivial. Here, at the level of LFP, we observed similarities between species in the interaction between ongoing and evoked activity: the largest evoked responses tended to be preceded by positive deflection in the LFP, and the smallest evoked responses were preceded by negative deflection in the LFP. This relationship, the negative interaction phenomenon, points to a non-additive interaction between ongoing and evoked activity and is also observed in both invasive and non-invasive recordings in humans (32,55,59,60). Establishing parallels between cortical dynamics on a well-defined task, such as sensory detection, between humans and animal models is an important direction for future studies.

In summary, we have developed a framework for the prediction of highly variable, singletrial sensory-evoked responses and shown that this prediction based on cortical state classification can be used to enhance the readout of sensory inputs. Utilizing state-dependent decoders for brain-computer interfaces has been shown to greatly improve the readout of motor commands from cortical activity (61,62), at the very end-stage of cortical processing. We suggest this natural extension of signal detection theory shows how to solve a problem that the brain faces at each stage of processing: how to adaptively read out a signal from a dynamical system constantly generating its own internal activity.

## Methods

### Animal preparation, surgery and habituation to head fixation

All procedures were approved by the 1nstitutional Animal Care and Use Committee at the Georgia 1nstitute of Technology (Protocol Number A16104) and were in agreement with guidelines established by the National 1nstitutes of Health. Six nine to twenty-six week old male C57BL/6J mice were used in this study. Mice were maintained under 1-2% isoflurane anesthesia while being implanted with a custom-made head-holder and a recording chamber. The location of the barrel column targeted for recording was functionally identified through intrinsic signal optical imaging (1SO1) under 0.5-1% isoflurane anesthesia. Recordings were targeted to B1, B2, C1, C2, and D2 barrel columns. Mice were habituated to head fixation, paw restraint and whisker stimulation for 3-7 days before proceeding to electrophysiological recordings. Following termination of the recordings, animals were anesthetized (isoflurane, 4-5%, for induction, followed by a euthanasia cocktail injection) and perfused.

### Electrophysiology

Local field potential was recorded using silicon probes (A1×32-5mm-25-177, NeuroNexus, USA) with 32 recording sites along a single shank covering 775 µm in depth. The probe was coated with Di1 (1,1’-dioctadecyl-3,3,3’3’-tetramethylindocarbocyanine perchlorate, 1nvitrogen, USA) for post hoc identification of the recording site. The probe contacts were coated with a PEDOT polymer (63) to increase signal-to-noise ratio. Contact impedance measured between 0.3 MOhm and 0.7 MOhm. The probe was inserted with a 35° angle relative to the vertical, until a depth of about 1000 µm. Continuous signals were acquired using a Cerebus acquisition system (Blackrock Microsystems, USA). Signals were amplified, filtered between 0.3 Hz and 7.5 kHz and digitized at 30 kHz.

### Whisker stimulus

Mechanical stimulation was delivered to a single contralateral whisker corresponding to the barrel column identified through 1SO1 using a galvo motor (Cambridge Technologies, USA). The galvo motor was controlled with millisecond precision using a custom software written in Matlab (Mathworks, USA). The whisker stimulus followed a sawtooth waveform (16 ms duration) of various velocities (1000 deg/s, 500 deg/s, 250 deg/s, 100 deg/s) delivered in the caudo-rostral direction. To generate stimuli of different velocity, the amplitude of the stimulus was changed while its duration remained fixed. Whisker stimuli of different velocities were randomly presented in blocks of 21 stimuli, with a pseudo-random inter-stimulus interval of 2 to 3 seconds and an inter-block interval of a minimum of 20 seconds. The total number of whisker stimuli across all velocities presented during a recording session ranged from 196 to 616 stimuli.

### LFP pre-processing and channel quality criteria

For analysis, the LFP was down-sampled to 2 kHz. The LFP signal entering the processing pipeline is raw, with no filtering beyond the anti-aliasing filters used at acquisition, enabling future use of these methods for real-time control. Prior to the analysis, signal quality on each channel was verified. We analyzed the power spectrum of LFP recorded on each channel for line noise at 60 Hz. In some cases, line noise could be mitigated by fitting the phase and amplitude of a 60-Hz sinusoid, as well as harmonics up to 300 Hz, over a 500-ms period in the pre-stimulus epoch, then extrapolating the sinusoid over the stimulus window and subtracting. A small number of channels displayed slow, irregular drift (2 or 3 of 32 channels) and these were discarded. All other channels were used.

### CSD calculation

Current source density (CSD) analysis was used for two different purposes: first, to functionally determine layers based on the average stimulus-evoked response, and second, to analyze the pre-stimulus activity (in single trials) to localize sinks and sources generating the predictive signal. We describe the general method used here. Prior to computing the current source density (CSD), each channel was scaled by its standard deviation to normalize impedance variation between electrodes. We then implemented the kernel CSD method (Potworowski et al 2012) to compute CSD on single trials. This method was chosen because it accommodates irregular spacings between electrodes, which occurs when recordings on a particular contact do not meet quality standards outlined above. To determine the best values for the kernel method parameters (regularization parameter, *λ*; source extent in x-y plane, *r*; and source extent in z-plane, *R*) we followed the suggestion of Potworowski (2012) and selected the parameter choices that minimize error in the reconstruction of LFP from the CSD. These parameters were similar across recordings, so for all recordings we used: *λ* = 0.0316; *r* = 200µ*m; R* = 37.5µ*m*.

### Functional assignment of cortical layers

The trial-averaged evoked response was computed on each trial by subtracting the pre-stimulus baseline (average over 200 ms prior to stimulus delivery) and computing the average across trials. The CSD of this response profile was computed as described above. The center of layer 4 was determined by finding the largest peak of the trial-averaged evoked LFP response as well as the location of the first, large sink in the trial-averaged sensory-evoked CSD response. We assume a width of 205 µm for layer 4, based on published values for mice (Hooks et al 2011).

### Analysis framework

#### Defining features of ongoing LFP

The ongoing LFP activity was characterized over the 2-s prestimulus window using “activation” and “power ratio.” LFP activation is defined here as the LFP averaged over the 10 ms pre-stimulus relative to the baseline value of the LFP averaged over 1000 ms to -200 ms. We observed that the estimate of pre-stimulus phase was biased by filter leakage from the evoked response (if the evoked response was not truncated) and biased by truncation otherwise. This filter leakage causes a spurious increase in classifier performance, because it reflected not only the evoked response but also the single-trial variability in the evoked response, which was easily read out by the classifier. Therefore, we used the simpler measure of LFP activation in place of pre-stimulus LFP phase. For the weighted average multi-channel LFP activation (Fig. 4), we performed a spatial PCA during the spontaneous activity and retained up to 3 spatial modes, or the number of modes that explained >95% of the variance, whichever was fewer. We then projected the pre-stimulus multi-channel LFP into spatial PC space. The power spectral density (PSD) was estimated over the 2-s pre-stimulus period on single trials using the fast Fourier transform. The power ratio was computed from the PSD curve as the area over the “L” range (1-5 Hz, or variable) normalized by the area over the “W” range (1-50 Hz). Analyses were also carried out using a 2-s pre-stimulus window and multi-taper PSD estimate (with NW = 2, 3 tapers), and the results were qualitatively unchanged.

#### Parameterizing the variable sensory-evoked LFP responses

Evoked responses were quantified using PCA to determine for each channel a small set of temporal basis functions that enabled us to parameterize the evoked response over the first 25 ms following sensory stimulation. The most prominent features in the PCA basis reflect the important features that one might define manually (amplitude and latency/duration of the cortical response; Fig. 2).

An alternative to the PCA representation of the evoked response is to attempt to directly measure identified features, such as peak response amplitude and response latency or duration (64-66). At the level of single trials, such metrics are susceptible to distortion from highfrequency fluctuations and require substantial pre-processing (64). This is avoided by using the linear projection of the evoked response onto the PCs. Further, we have focused on the 25-ms period post-stimulus, and most of the methods developed around templates matching or differential variable components are better suited for slower signals (66).

We also computed the full (32-channel) spatio-temporal PCA basis. To compute the full spatial PCA, we used a cross-validation method in which 20% of observations were randomly dropped and we utilized an alternating least squares approach to find coefficients and weights. A small number (2-3) of full spatiotemporal modes captures captured >90% of the response variability. Moreover, the weights obtained for the layer-4 PCs were nearly identical to the full spatiotemporal mode weights, indicating that variability in the LFP was dominated by layer 4 variability and justifying our focus on predicting layer 4 variability alone.

#### Mathematical formalism

We represent the discretely sampled (2 kHz) LFP time series by *x(t)*. The time series of the evoked LFP response to a stimulus at time *t*_*i*_ (in milliseconds) in the subsequent 25-ms period is represented as a length-*N*_*E*_ vector 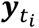, built up from *x(t)* (Δ *t* = 0.5 ms, *N*_*E*_ = 50):

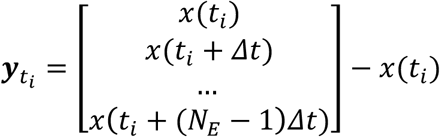

Note that the LFP at the time of stimulus delivery is subtracted in order to return the evoked response 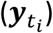 The average evoked response over *N*_*T*_ trials is ***ξ***_0_:

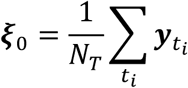

and the single-trial response is written as the sum of the mean response and a noise term ***η*:**

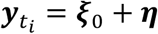

Here, we expand the evoked response at time *t*_*i*_ in terms of its first *N*_*C*_ components across trials:

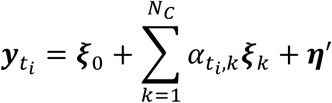

where the vectors ***ξ***_*k*_ are obtained from the principal components of evoked responses 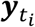, i.e. the eigenvectors of *YY*^*T*^, where the *i*^*th*^ column of *Y* is 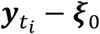.

#### Fitting the state classifier

Here, we are motivated to find the pre-stimulus features that best predict single-trial evoked responses, and to then define “cortical state” based on those combinations of features that produce similar cortical responses. Therefore, we aim to find some function of the pre-stimulus features that generates a prediction of the coefficients 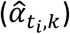 The predicted response for a stimulus at time *t*_*i*_ is 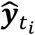:

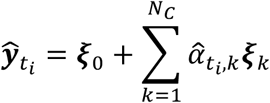

We considered the first two components, so *N*_*C*_ = 2. Evoked response weights 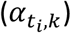 were split into equipartitioned groups based on quintiles of the weight distribution. Taking as predictors (inputs to the classifier) the features of pre-stimulus activity described above, the classifier was trained to predict to which group the evoked response belonged. Each evoked mode (*k*) is independently predicted.

Each recording session was divided into a training/cross-validation set (70% of trials) and a test set (30% of trials). Classifiers were fit on the training/CV set using five-fold cross-validation, and predictions and accuracy were recorded for the cross-validation trials. Optimized parameters, such as ranges for the L range of the power ratio, were selected on the basis of error in the crossvalidation set. Reported classifier performance (fraction of variance explained, fVE, see below) was then calculated on the reserved test set. For the Gaussian-kernel SVM, we used a medium-scale kernel (3; predictors range from 0 to 1 (power raito) and +/1 (mV, activation)) and box constraint of 1. Results were not sensitive to this choice. We also tested linear kernel SVMs as well as more complicated decoders, such as k-nearest neighbor classifiers, but did not find any increase in performance across recordings (not shown). All classifiers were fit using built-in Matlab routines, fitcecoc, from the classification toolbox (MATLAB 2017a, Mathworks, Natick, MA).

#### Classifier performance quantification (fVE)

Fraction of variance explained (fVE) is defined as the normalized difference between the total trial-by-trial variance (summed over time) and the trial-averaged squared error between the predicted response and the data (also summed over time):

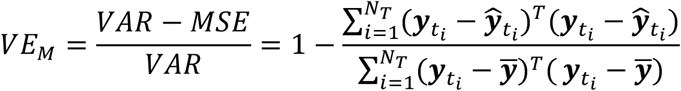

#### Shuffled data fVE

trial labels (i.e. the quintile of the PC distribution to which a trial was assigned) were shuffled. The classifier (fit on unshuffled data) was applied to the pre-stimulus activity, and fVE was computed for the predicted classes supposing that the shuffled labels were accurate.

#### Uncertainty in fVE from finite quantity of data

Differences in fVE between classifiers arise from differences in the accuracy of each classifier in predicting a subset of reserved data and therefore partially reflect the true accuracy as well as random effects arising from the finite quantity of data. We use a resampling approach to estimate the uncertainty in fVE. Because the test set is stratified (i.e. it is balanced across labels (quintiles)), we use the leave-one-out jackknife: drop five trials (one from each quintile of the response), compute the fVE, then repeat until each trial has been dropped. The sample standard deviation of the resampled fVE distribution is reported as the uncertainty in the fVE estimate.

### State-blind and state-aware detection

The matched filter ideal observer analysis (37) is implemented as follows. The *score s(t)* is constructed by taking the dot product of the evoked responses ***y***_*t*_ with a filter matched to the average evoked response:

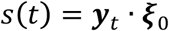

This is equivalent to computing the sum

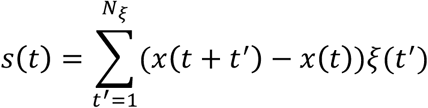

In the standard encoding model, if ***η*** is zero-mean white noise, this gives a signal distribution

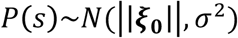

where 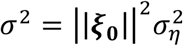 and a noise distribution with mean 0. 1n practice, we do not parameterize the distribution, because ***η*** is not uncorrelated white noise, and work from the score distribution directly.

For the state-aware decoder, we use the prediction 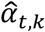 of evoked responses

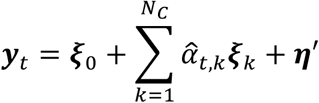

This changes the score to

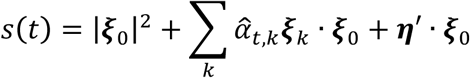

Typically, one of the first two PCs (***ξ***_1_ or ***ξ***_2_) has a very similar shape to ***ξ***_0_, while the other one has both positive and negative components (Fig. 2, Suppl. Fig. 1-2). For the state-aware threshold, we use state predictions for the component that is more similar to ***ξ***_0_, as indicated in Supplemental Figure 1-2.

An event is detected at time *t* for threshold *θ* when s (*t*)*> θ* is a local maximum that is separated from the nearest peak by at least 15 ms and has a minimum prominence (i.e. drop in *s* before encountering another peak that was higher than the original peak) of **|*ξ*** _0_ **|^2^/2.**

### Data availability statement

The datasets generated during and analyzed for this study will be made available in the Dryad data repository.

## Supplemental Figures

**Supplemental Figure 1.**
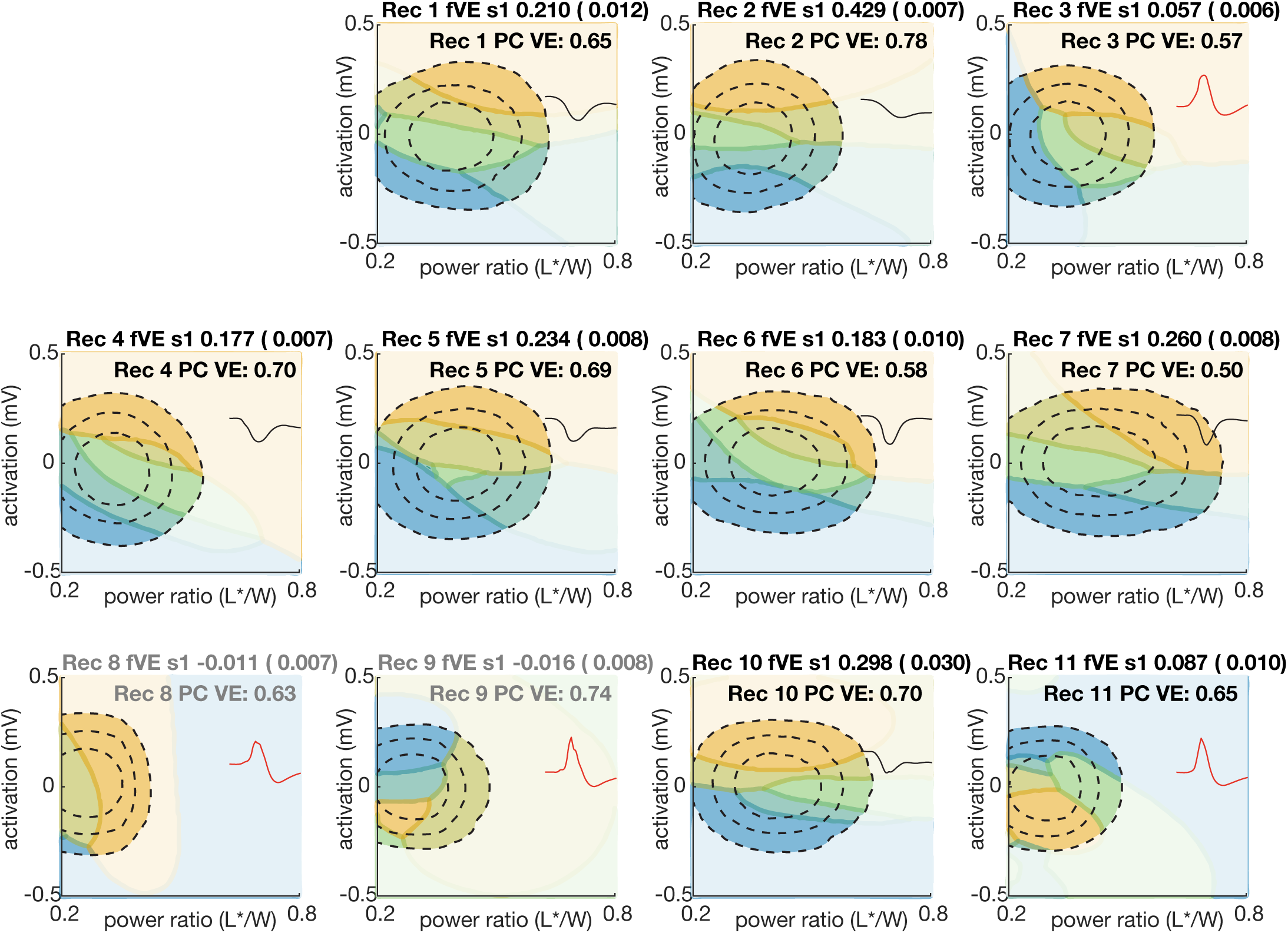
(associated with Fig. 2): Classifier boundaries for the prediction of the weight onto the first principal component of the evoked responses for recordings 1 to 11. Each panel is a recording. Fraction of variance explained by the classifier (“fVE s1”) is in the title of each panel (bootstrapped SE in parentheses). Gray indicates that the classifier did not predict above chance levels (shuffle test). PC VE is the variance explained for the first principal component for each recording. (fVE cannot exceed PC VE.) Inset shape shows the principal component (PC) loadings. Black is “amplitude” (Rec. 1, 2, 4, 5, 6, 7, 10) and red is “sharpness/latency” (Rec. 3, 8, 9, 11). Compare to Supplemental Figure 2, which shows the same plots for the second principal component.

**Supplemental Figure 2.**
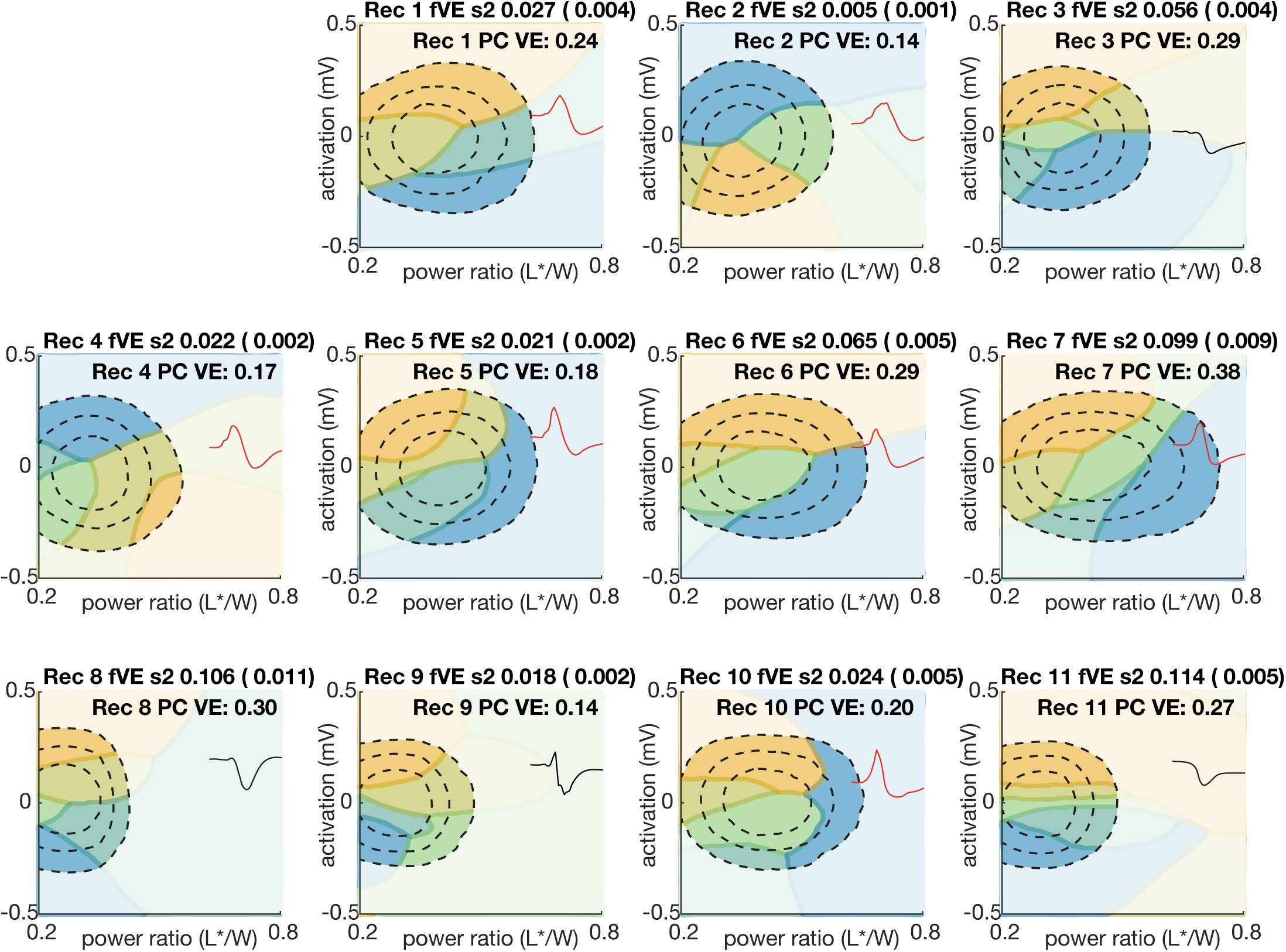
(associated with Fig. 2): Classifier boundaries for the prediction of the weight onto the second principal component of the evoked responses for recordings 1 to 11. Each panel is a recording. Fraction of variance explained by the classifier (fVE s2) is in the title of each panel (bootstrapped SE in parentheses). PC VE is the variance explained for the second principal component for each recording. (fVE cannot exceed PC VE.) Gray indicates that the classifier did not predict above chance levels (shuffle test). Inset shape shows the principal component (PC) loadings. Black is “amplitude” (Rec. 3, 8, 9, 11) and red is “sharpness/latency” (Rec. 1, 2, 4, 5, 6, 7, 10). Compare to Supplemental Figure 1, which shows the same plots for the first principal component.

**Supplemental Figure 3.**
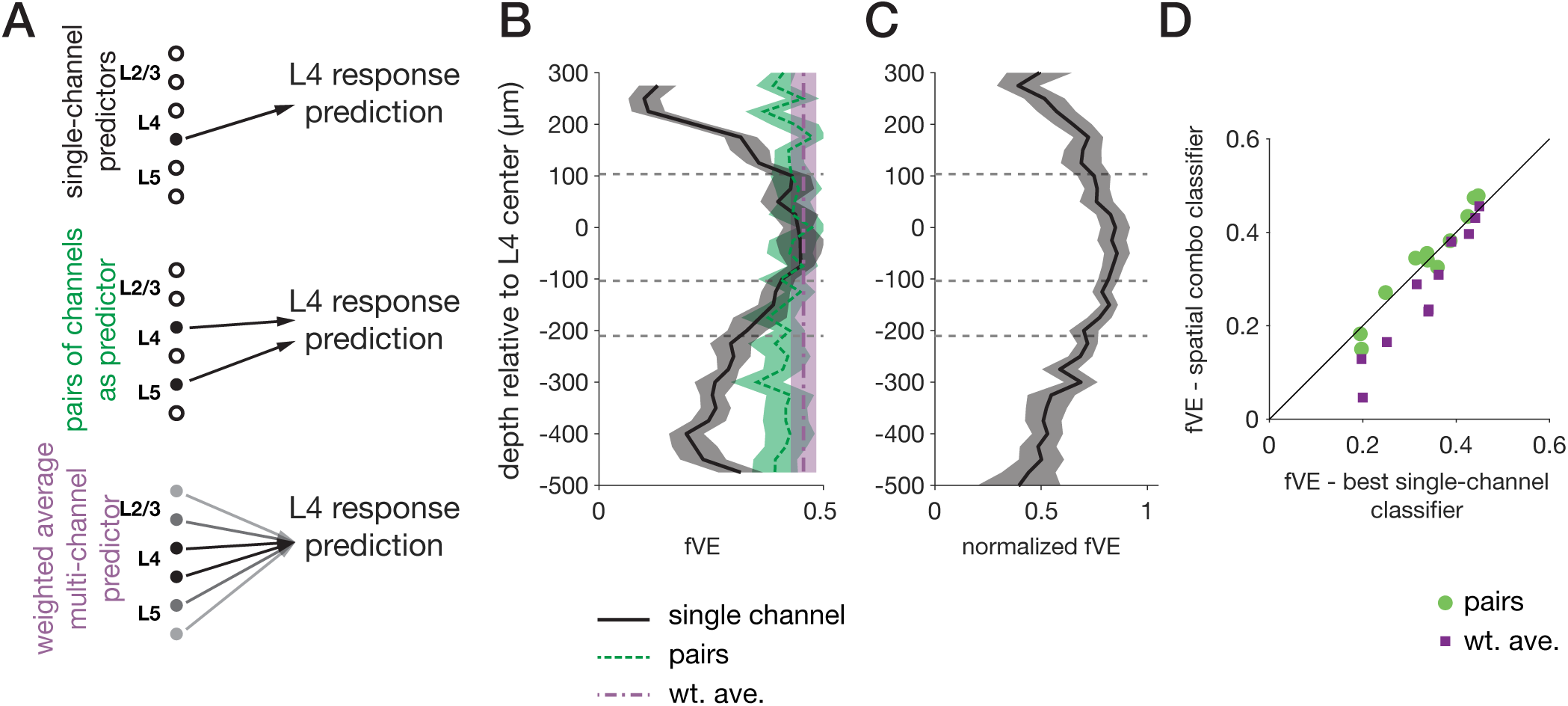
(associated with Figure 4). Comparison of performance for more complex classifiers. A-D: Same as Fig. 4, but including a “multi-channel predictor,” constructed from the projection of pre-stimulus activity onto the top spatial modes in spontaneous LFP determined by PCA.

**Supplemental Figure 4.**
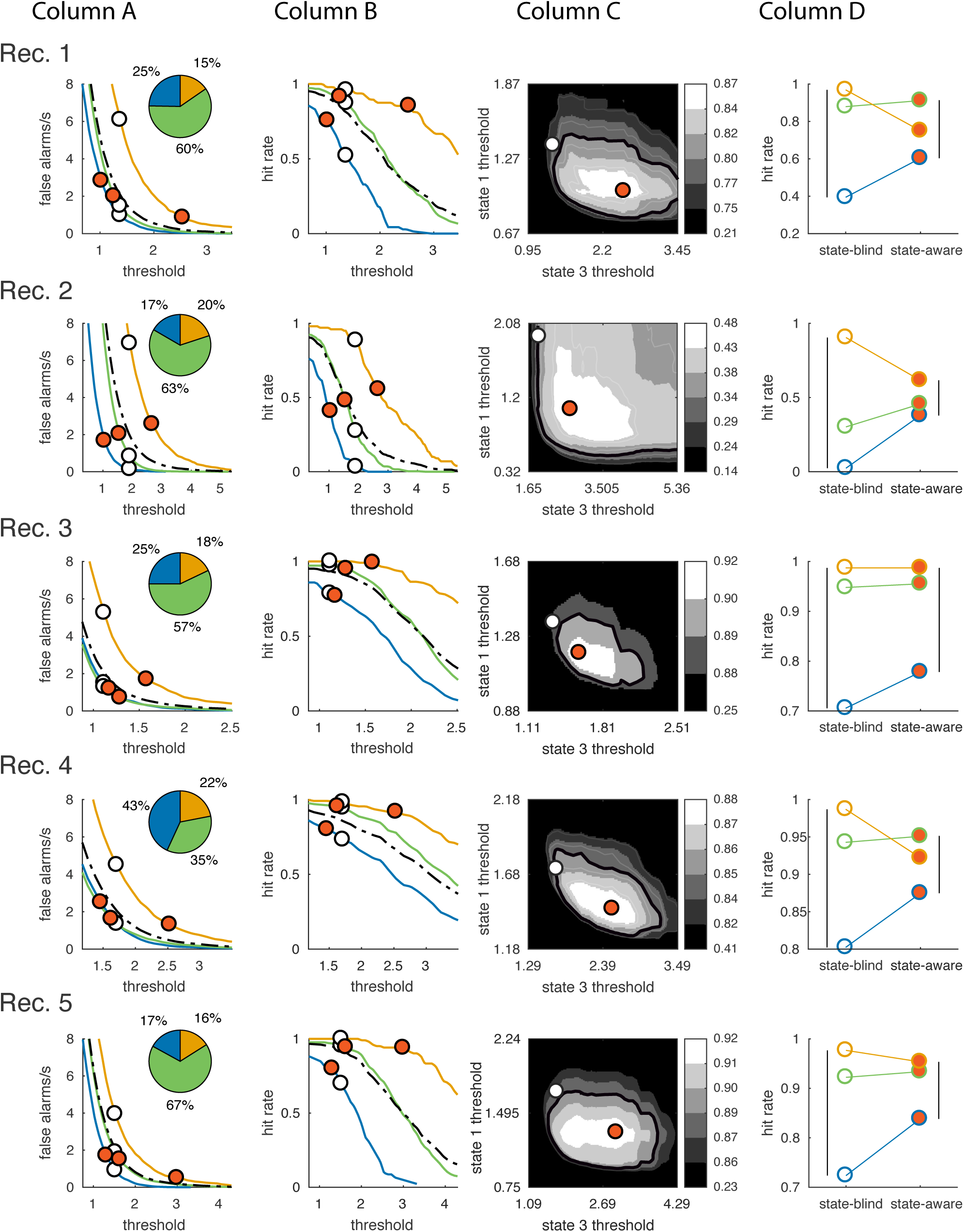
(associated with Figure 6): Optimized thresholds for the state-aware observer (panels A-D of Figure 6) for recordings 1 through 5. Column A: false alarms per s vs. threshold. Inset: fraction of spontaneous activity in each state. Column B: hit rates vs threshold. Column C: State-aware detection rates at fixed false alarm rate for combinations of state1 and state3 thresholds. Red contour is the detection rate with fixed threshold. Column D: Hit rate, conditioned on pre-stimulus state, in the state-blind and state-aware cases.

**Supplemental Figure 5.**
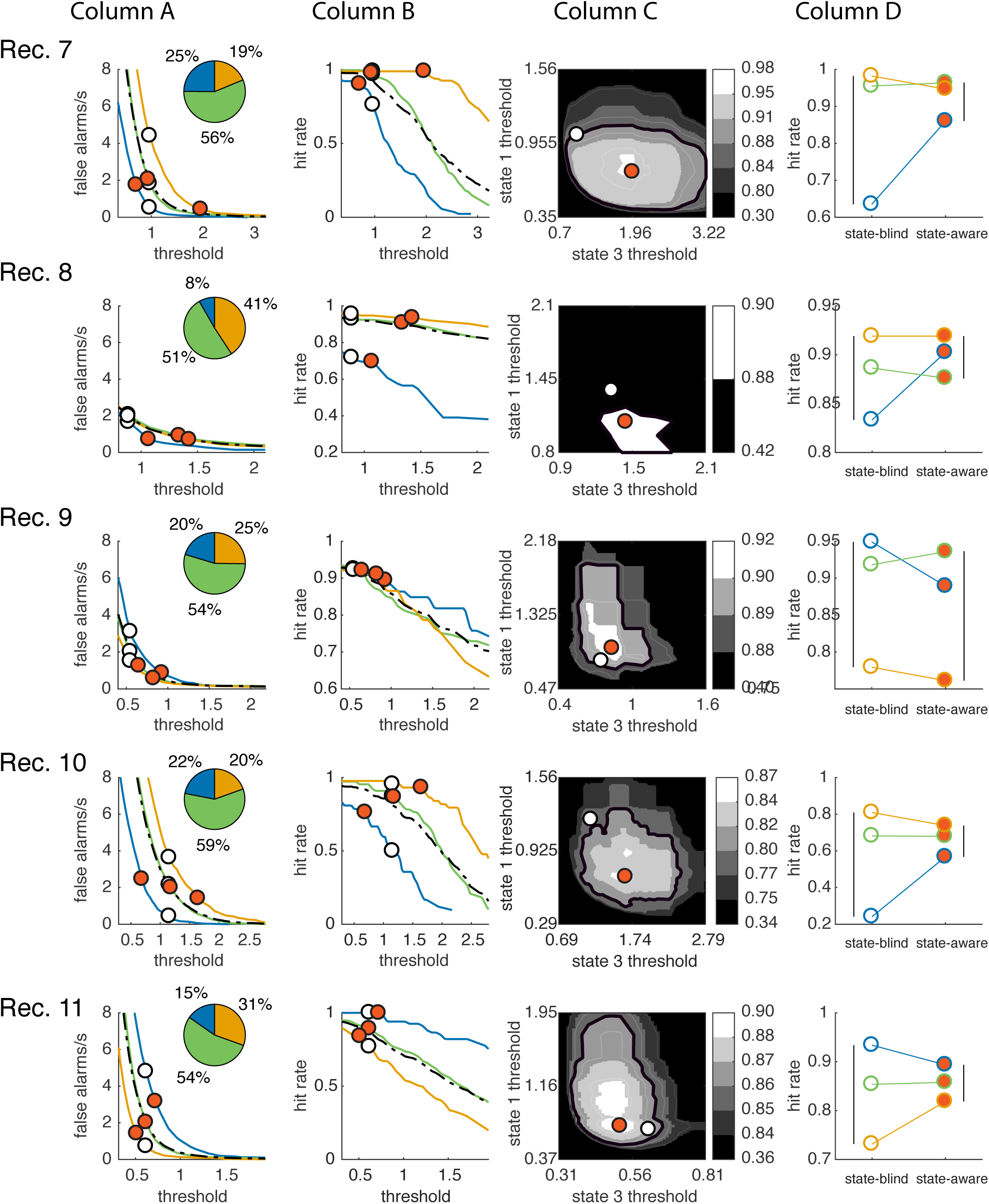
(associated with Figure 6): Optimized thresholds for the state-aware observer (panels A-D of Figure 6) for recordings 7 through 11. Column A: false alarms per s vs. threshold. Inset: fraction of spontaneous activity in each state. Column B: hit rates vs threshold. Column C: State-aware detection rates at fixed false alarm rate for combinations of state1 and state3 thresholds. Red contour is the detection rate with fixed threshold. Column D: Hit rate, conditioned on pre-stimulus state, in the state-blind and state-aware cases. Recording 6 is shown in Figure 6.

## References

1. Vogels R, Spileers W, Orban GA. The response variability of striate cortical neurons in the behaving monkey. Exp Brain Res. 1989;77(2):432–6.

2. Kara P, Reinagel P, Reid RC. Low Response Variability in Simultaneously Recorded Retinal, Thalamic, and Cortical Neurons. 2000;27:635–46.

3. Gur M, Snodderly DM. High response reliability of neurons in primary visual cortex (V1) of alert, trained monkeys. Cereb Cortex. 2006;16(6):888–95.

4. Ma WJ, Beck JM, Latham PE, Pouget A. Bayesian inference with probabilistic population codes. 2006;9(11):1432–8.

5. Orbán G, Berkes P, Fiser J, Lengyel M. Neural Variability and Sampling-Based Probabilistic Representations in the Visual Cortex. Neuron. 2016 Oct;92(2):530–43.

6. Arazi A, Censor N, Dinstein I. Neural Variability Quenching Predicts Individual Perceptual Abilities. J Neurosci. 2017;37(1):97–109.

7. Arieli A, Sterkin A, Grinvald A, Aertsen A. Dynamics of Ongoing Activity: Explanation of the Large Variability in Evoked Cortical Responses. Science (80-). American Association for the Advancement of Science; 1996 Sep;273(5283):1868–71.

8. Petersen CCH, Hahn TTG, Mehta MR, Grinvald A, Sakmann B. Interaction of sensory responses with spontaneous depolarization in layer 2/3 barrel cortex. Proc Natl Acad Sci U S A. 2003;100(23):13638–43.

9. Haslinger R, Ulbert I, Moore CI, Brown EN, Devor A. Analysis of LFP phase predicts sensory response of barrel cortex. J Neurophysiol. 2006;96(3):1658–63.

10. Haider B, Duque A, Hasenstaub A, Yu Y, McCormick D a. Enhancement of visual responsiveness by spontaneous local network activity in vivo. J Neurophysiol. 2007;97(6):4186–202.

11. Marguet SL, Harris KD. State-Dependent Representation of Amplitude-Modulated Noise Stimuli in Rat Auditory Cortex. 2011;31(17):6414–20.

12. Pachitariu M, Lyamzin DR, Sahani M, Lesica N a. State-dependent population coding in primary auditory cortex. J Neurosci. 2015;35(5):2058–73.

13. Reig R, Zerlaut Y, Vergara R, Destexhe A, Sanchez-Vives M V. Gain Modulation of Synaptic Inputs by Network State in Auditory Cortex In Vivo. J Neurosci. 2015;35(6):2689–702.

14. Curto C, Sakata S, Marguet S, Itskov V, Harris KD. A simple model of cortical dynamics explains variability and state-dependence of sensory responses in urethane-anesthetized auditory cortex. J Neurosci. 2009;29(34):10600–12.

15. Gutnisky DA, Beaman CB, Lew SE, Dragoi V. Spontaneous Fluctuations in Visual Cortical Responses Influence Population Coding Accuracy. 2017;(February):1409–27.

16. Baria AT, Maniscalco B, He BJ. Initial-state-dependent, robust, transient neural dynamics encode conscious visual perception. PLoS Comput Biol. 2017;13(11):1–29.

17. Monto S, Palva S, Voipio J, Palva JM. Very Slow EEG Fluctuations Predict the Dynamics of Stimulus Detection and Oscillation Amplitudes in Humans. J Neurosci. 2008;28(33):8268–72.

18. Ress D, Heeger DJ. Neuronal correlates of perception in early visual cortex. Nat Neurosci. 2003;6(4):414–20.

19. Fox MD, Snyder AZ, Zacks JM, Raichle ME. Coherent spontaneous activity accounts for trial-to-trial variability in human evoked brain responses. Nat Neurosci. 2006;9(1):23–5.

20. Boly M, Balteau E, Schnakers C, Degueldre C, Moonen G, Luxen A, et al. Baseline brain activity fluctuations predict somatosensory perception in humans. Proc Natl Acad Sci. 2007 Jul 17;104(29):12187–92.

21. Li Q, Hill Z, He BJ. Spatiotemporal Dissociation of Brain Activity Underlying Subjective Awareness, Objective Performance and Confidence. J Neurosci. 2014;34(12):4382–95.

22. De Lafuente V, Romo R. Neuronal correlates of subjective sensory experience. Nat Neurosci. 2005;8(12):1698–703.

23. van Vugt B, Dagnino B, Vartak D, Safaai H, Panzeri S, Dehaene S, et al. The threshold for conscious report: Signal loss and response bias in visual and frontal cortex. Science (80-). 2018;7186(March).

24. Kyriakatos A, Petersen CCH, Kyriakatos A, Sadashivaiah V, Zhang Y, Motta A, et al. Voltage-sensitive dye imaging of mouse neocortex during a whisker detection task during a whisker detection task. 2017;4(3).

25. Waiblinger C, Whitmire CJ, Sederberg A, Stanley GB, Schwarz C. Primary tactile thalamus spiking reflects cognitive signals. J Neurosci. 2018;38(21).

26. Sachidhanandam S, Sreenivasan V, Kyriakatos A, Kremer Y, Petersen CCH. Membrane potential correlates of sensory perception in mouse barrel cortex. Nat Neurosci. 2013;16(11):1671–7.

27. Green DM, Swets JA. Signal Detection Theory and Psychophysics. New York: John Wiley and Sons; 1966.

28. Wang Q, Webber RM, Stanley GB. Thalamic synchrony and the adaptive gating of information flow to cortex. Nat Neurosci. Nature Publishing Group; 2010;13(12):1534–41.

29. Ritt JT, Andermann ML, Moore CI. Embodied Information Processing: Vibrissa Mechanics and Texture Features Shape Micromotions in Actively Sensing Rats. Neuron. 2008;57(4):599–613.

30. Arabzadeh E, Zorzin E, Diamond ME. Neuronal encoding of texture in the whisker sensory pathway. PLoS Biol. 2005;3(1).

31. Hooks BM, Hires SA, Zhang YX, Huber D, Petreanu L, Svoboda K, et al. Laminar analysis of excitatory local circuits in vibrissal motor and sensory cortical areas. PLoS Biol. 2011;9(1).

32. He BJ. Spontaneous and Task-Evoked Brain Activity Negatively Interact. 2013;33(11):4672–82.

33. Nicholson C, Freeman JA. Theory of current source-density analysis and determination of conductivity tensor for anuran cerebellum. J Neurophysiol. 1975;38(2).

34. Mitzdorf U. Current source-density method and application in cat cerebral cortex: investigation of evoked potentials and EEG phenomena. Physiol Rev. 1985;65(1):37–100.

35. Pettersen KH, Devor A, Ulbert I, Dale AM, Einevoll GT. Current-source density estimation based on inversion of electrostatic forward solution: Effects of finite extent of neuronal activity and conductivity discontinuities. J Neurosci Methods. 2006 Jun;154(1–2):116–33.

36. Potworowski J, Jakuczun W, Łȩski S, Wójcik DK. Kernel Current Source Density Method. Neural Comput. 2012;24(2):541–75.

37. Oppenheim A, Verghese G. Signals, Systems and Inference, Chapter 14: Signal Detection. 2010;247–62.

38. Poulet JFA, Petersen CCH. Internal brain state regulates membrane potential synchrony in barrel cortex of behaving mice. 2008;454(August).

39. Fernandez LMJ, Comte J, Le Merre P, Lin J-S, Salin P-A, Crochet S. Highly Dynamic Spatiotemporal Organization of Low-Frequency Activities During Behavioral States in the Mouse Cerebral Cortex. Cereb Cortex. 2016 Oct 14;(January 2018):5444–62.

40. Olcese U, Oude Lohuis MN, Pennartz CMA. Sensory Processing Across Conscious and Nonconscious Brain States: From Single Neurons to Distributed Networks for Inferential Representation. Front Syst Neurosci. 2018;12(October):49.

41. McGinley MJ, Vinck M, Reimer J, Batista-Brito R, Zagha E, Cadwell CR, et al. Waking State: Rapid Variations Modulate Neural and Behavioral Responses. Neuron. Elsevier Inc.; 2015 Sep;87(6):1143–61.

42. Gervasoni D, Lin S-C, Ribeiro S, Soares ES, Pantoja J, Nicolelis MAL. Global Forebrain Dynamics Predict Rat Behavioral States and Their Transitions. J Neurosci. 2004 Dec 8;24(49):11137–47.

43. Steriade M. Corticothalamic resonance, states of vigilance and mentation. Neuroscience. 2000;101(2):243–76.

44. Vinck M, Batista-Brito R, Knoblich U, Cardin JA. Arousal and Locomotion Make Distinct Contributions to Cortical Activity Patterns and Visual Encoding. Neuron. Elsevier Inc.; 2015;86(3):740–54.

45. Pesaran B, Vinck M, Einevoll GT, Sirota A, Fries P, Siegel M. Investigating large-scale brain dynamics using field potential recordings: Analysis and interpretation. Nat Neurosci. Springer US; 2018;

46. Einevoll GT, Pettersen KH, Devor A, Ulbert I, Halgren E, Dale AM. Laminar population analysis: estimating firing rates and evoked synaptic activity from multielectrode recordings in rat barrel cortex. J Neurophysiol. 2007;97(3):2174–90.

47. Reyes-Puerta V, Sun J-J, Kim S, Kilb W, Luhmann HJ. Laminar and Columnar Structure of Sensory-Evoked Multineuronal Spike Sequences in Adult Rat Barrel Cortex In Vivo. Cereb Cortex. 2015;25(8):2001–21.

48. Yang H, Kwon SE, Severson KS, O’Connor DH. Origins of choice-related activity in mouse somatosensory cortex. Nat Neurosci. 2015;19(1):127–34.

49. Nienborg H, R. Cohen M, Cumming BG. Decision-Related Activity in Sensory Neurons: Correlations Among Neurons and with Behavior. Annu Rev Neurosci. 2012 Jul 21;35(1):463–83.

50. Montijn XJS, Olcese U, Pennartz XCMA. Visual Stimulus Detection Correlates with the Consistency of Temporal Sequences within Stereotyped Events of V1 Neuronal Population Activity. 2016;36(33):8624–40.

51. Palmer C, Cheng S-Y, Seidemann E. Linking Neuronal and Behavioral Performance in a Reaction-Time Visual Detection Task. J Neurosci. 2007;27(30):8122–37.

52. Hong YK, Lacefield CO, Rodgers CC, Bruno RM. Sensation, movement and learning in the absence of barrel cortex. Nature. Springer US; 2018;561(7724):542–6.

53. O’Connor DH, Peron SP, Huber D, Svoboda K. Neural activity in barrel cortex underlying vibrissa-based object localization in mice. Neuron. Elsevier Inc.; 2010;67(6):1048–61.

54. Stüttgen MC, Schwarz C. Barrel cortex: What is it good for? Neuroscience. 2018;368:3–16.

55. Sadaghiani S, Hesselmann G, Kleinschmidt A. Distributed and Antagonistic Contributions of Ongoing Activity Fluctuations to Auditory Stimulus Detection. J Neurosci. 2009;29(42):13410–7.

56. Iemi L, Busch NA. Moment-to-Moment Fluctuations in Neuronal Excitability Bias Subjective Perception Rather than Strategic Decision-Making. Eneuro. 2018;5(3):ENEURO.0430-17.2018.

57. Busch NA, Dubois J, VanRullen R. The Phase of Ongoing EEG Oscillations Predicts Visual Perception. J Neurosci. 2009 Jun 17;29(24):7869–76.

58. Iemi XL, Chaumon M, Se X, Crouzet M, Busch NA. Spontaneous Neural Oscillations Bias Perception by Modulating Baseline Excitability. 2017;37(4):807–19.

59. He BJ, Zempel JM. Average Is Optimal: An Inverted-U Relationship between Trial-to-Trial Brain Activity and Behavioral Performance. PLoS Comput Biol. 2013;9(11).

60. Hesselmann G, Kell CA, Eger E, Kleinschmidt A. Spontaneous local variations in ongoing neural activity bias perceptual decisions. Proc Natl Acad Sci. 2008;105(31):10984–9.

61. Aggarwal V, Mollazadeh M, Davidson AG, Schieber MH, Thakor N V. State-based decoding of hand and finger kinematics using neuronal ensemble and LFP activity during dexterous reach-to-grasp movements. J Neurophysiol. 2013;109(12):3067–81.

62. Pandarinath C, Gilja V, Blabe CH, Nuyujukian P, Sarma AA, Sorice BL, et al. Neural population dynamics in human motor cortex during movements in people with ALS. ELife. 2015;4:e07436.

63. Ludwig KA, Uram JD, Yang J, Martin DC, Kipke DR. Chronic neural recordings using silicon microelectrode arrays electrochemically deposited with a poly(3,4- ethylenedioxythiophene) (PEDOT) film. J Neural Eng. 2006;3(1):59–70.

64. Ahmadi M, Quian Quiroga R. Automatic denoising of single-trial evoked potentials. Neuroimage. Elsevier Inc.; 2013;66:672–80.

65. Mahmud M, Vassanelli S. Processing and analysis of multichannel extracellular neuronal signals: State-of-the-art and challenges. Front Neurosci. 2016;10(JUN):1–12.

66. Truccolo W, Knuth KH, Shah A, Bressler SL, Schroeder CE, Ding M. Estimation of single-trial multicomponent ERPs: Differentially variable component analysis (dVCA). Biol Cybern. 2003;89(6):426–38.

